# Strawberry sweetness and consumer preference are enhanced by specific volatile compounds

**DOI:** 10.1101/2020.12.04.410654

**Authors:** Zhen Fan, Tomas Hasing, Timothy S. Johnson, Drake M. Garner, Christopher R. Barbey, Thomas A. Colquhoun, Charles A. Sims, Marcio F. R. Resende, Vance M. Whitaker

## Abstract

Breeding crops for improved flavor is challenging due to the high cost of sensory evaluation and the difficulty of connecting sensory experience to chemical composition. The main goal of this study was to identify the chemical drivers of sweetness and consumer liking for fresh strawberries (*Fragaria ×ananassa*). Fruit of 148 strawberry samples from cultivars and breeding selections were grown and harvested over seven years and were subjected to both sensory and chemical analyses. Each panel consisted of at least 100 consumers, resulting in more than 15,000 sensory data points per descriptor. Three sugars, two acids and 113 volatile compounds were quantified. Consumer liking was highly associated with sweetness intensity, texture liking, and flavor intensity, but not sourness intensity. Partial least square analyses revealed 20 volatile compounds that increased sweetness perception independently of sugars; 18 volatiles that increased liking independently of sugars; and 15 volatile compounds that had positive effects on both. Machine learning-based predictive models including sugars, acids, and volatiles explained at least 25% more variation in sweetness and liking than models accounting for sugars and acids only. Volatile compounds such as γ-dodecalactone; 5-hepten-2-one, 6-methyl; and multiple medium-chain fatty acid esters may serve as targets for breeding or quality control attributes for strawberry products. A genetic association study identified two loci controlling ester production, both on linkage group 6A. Co-segregating makers in these regions can be used for increasing multiple esters simultaneously. This study demonstrates a paradigm for improvement of fruit sweetness and flavor in which consumers drive the identification of the most important chemical targets, which in turn drives the discovery of genetic targets for marker-assisted breeding.

## Introduction

The cultivated strawberry (*Fragaria ×ananassa*) is one of the most widely grown fruit crops in the world due to its sweet and aromatic flavor and health-associated compounds including anthocyanins, antioxidants, fiber and ellagic acid^1^. Breeding for flavor in strawberry has been challenging due to the chemical complexity of the trait and the variability of the production environment. Targeted selection of flavor-associated chemicals in strawberry has thus far been restricted to increasing sugar content. In spite of improvements in eating quality over time, most strawberries on the market still do not meet consumer expectations of sweetness and flavor^2^. Efforts at flavor improvement must be reinforced by investigating the relationships between fruit chemical diversity and consumer preference.

During strawberry fruit ripening, changing auxin levels drive the accumulation of sugars and their derivatives, as well as secondary metabolites^3,4^. Sucrose, glucose, and fructose comprise the major soluble sugars in strawberry and are rapidly translocated from photosynthesizing organs to fruit, in synchrony with reductions in other sugars including xylose, galactose, sugar phosphates and sugar alcohols^3^. Besides sugars, amino acids, phenolic compounds and volatiles are the main indicators of fruit ripening. Strawberry aroma is determined by low-molecular-weight volatile compounds generated during ripening. Over 360 volatiles have been identified in the aroma of cultivated and wild strawberries, with about 280 of them reported in cultivated strawberry^5^. Esters, primarily methyl and ethyl esters, constitute 25-90% of the total abundance of volatiles in strawberry and provide fruity notes^6^. Lactones, cyclic esters providing aromas similar to those in peach, are prominent volatiles in some varieties^7^. Aldehydes such as 2-hexenal, (E)- and 3-hexenal, (Z)-contribute to green or fresh aromas^8^. Furanones such as furaneol and mesifurane are often associated with sweet aromas^9^.

Sensory analyses are necessary to comprehensively characterize fruit flavors, which are highly influenced by retronasal olfaction^10^. Odor Activity Values (OAVs), which are ratios of concentration to odor thresholds, were adopted in early studies to identify potent volatile compounds in strawberry^11^. These studies identified several compounds as important to strawberry flavor including (Z)-3-hexenal (green), 4-hydroxy-2,5-dimethyl-3(2H)-furanone (sweet), methyl butanoate (fruity), and ethyl butanoate (fruity). However, OAVs do not account for interactions between the matrix and the volatile, since the denominator only reflects odor intensity in a water solution^12^. Furthermore, OAVs are only based on orthonasal olfaction which is a different sensory experience than retronasal olfaction. Gas chromatography-olfactometry (GC-O) is another powerful tool to identify potent volatiles in food. Comparable results were obtained by GC-O and OAV for strawberry, agreeing on the most intense odorants^8,13^. However, while these studies identified potent volatiles, synergy among volatile compounds to produce human sensory responses and interactions between taste and retronasal olfaction were unexplored.

Studies on tomato flavor have highlighted the flaws of using OAVs exclusively for determining volatiles important to sensory perceptions^14^. Large-scale sensory and chemical studies have facilitated identification of flavor- or taste- associated volatiles^15^, allowing breeding efforts to focus on genetic improvement of a smaller set of important volatiles. Once natural genetic variations responsible for volatile biosynthesis within heirloom tomato populations were exploited, DNA marker-assisted breeding techniques were utilized to introgress desirable alleles for volatile biosynthesis genes into modern genetic backgrounds^16^. A recent study in blueberry identified genomic regions associated with those compounds and also explored correlations between sensory attributes and volatile compounds^17^. A previous study in strawberry by the present authors, utilizing approaches and techniques similar to Tieman *et al.*^15^, linked sugars, acids, and volatile data to consumer panel data for 35 strawberry genotypes^18^. Significant correlations between volatiles and sweetness and flavor intensities were found. Due to the two-year time-frame of the study, total sample size was limited to 54, and chemicals influencing consumer liking were not explored^18^. Marker-assisted selection in strawberry breeding for some flavor-associated volatiles has recently become a reality^19^. There is now a need to identify more sensory-associated volatile compounds in order to breed more comprehensively for strawberry flavor.

Genetic background is a major factor influencing primary and secondary metabolites in fresh strawberry fruit. Multiple studies have found that volatile compounds vary both qualitatively and quantitatively among strawberry varieties, leading to flavor differences^5–7,20^. Notable differences in volatile profiles have been observed in popular cultivars grown in the US, with more than ten-fold difference observed in all classes of volatiles^7,21,22^. Distinguishable separations between summer-grown European strawberry cultivars and two winter-grown Florida cultivars were found for 12 key strawberry volatiles^8^. During domestication, some desirable aroma-active volatiles have been lost over time^23^. Methyl cinnamate and methyl anthranilate, which are abundant in wild ancestor *F. vesca* and elicit pleasant aromas, are not detectable in modern cultivars^24^. Temperature is another critical factor that influences strawberry quality. In a single winter production season, temperature fluctuations drove substantial changes in fruit metabolic composition^25^. Cooler nights enhanced both sucrose accumulation and production of aroma compounds in a greenhouse setting^26^. Other environmental factors influencing strawberry fruit chemical abundance include light^27^, maturity, postharvest storage^6^, cultivation system^28^, and location^29^.

Recent studies have found that specific volatiles are capable of enhancing, not just flavor perception, but also sweetness perception in fresh fruit^15,18^. Hence, quantification of volatiles, sugars and acids should allow some degree of predictive ability for sensory characteristics, including sweetness. The recent development of machine learning models has facilitated prediction of sensory attributes and hedonic experiences based on chemistry. Partial least square (PLS) and its derivatives have been tested and validated for prediction of sensory attributes in different fruits, vegetables, and drinks^30–34^. In one example, artificial neural networks were used to simulate non-linear relationships for improvement of prediction accuracy for flavor intensity in blackcurrant^35^. To date, no predictive models for sensory characteristics or liking have been reported for fresh strawberries.

The main objectives of present study were to 1) explore the chemical drivers of consumer preference in fresh strawberries; 2) identify volatile compounds enhancing sweetness perception independently of sugars; and 3) predict sweetness perception and liking from chemical data. This work builds on previous research^18^, increasing the sample size approximately three-fold and applying updated mathematical and predictive models.

## Results

### Relationships among sensory and hedonic consumer responses

Over the course of seven years, 384 consumer panelists with an average age of 31.7 participated in evaluating strawberry samples. Panelists were 35% male and 65% female. Significant correlations were found among all pair-wise comparisons of sensory and hedonic ratings (Table S1A). Sweetness (r=0.714*), texture liking (r=0.783*) and flavor intensity (r=0.688*) were highly correlated with overall liking. Sourness (r=0.298*) had a significant but lower positive correlation with overall liking. Notably, sweetness had significant correlations with all other sensory ratings (r=0.411*, 0.592* and 0.838* for sourness, texture and flavor intensity, respectively). To remove the confounding effect of sweetness, partial correlations were calculated between liking and other attributes while controlling for sweetness (Table S1B). After controlling for sweetness, the correlation between sourness and liking reduced to an insignificant level (r=0.008), which was consistent with previous findings^18^. Conversely, this result was at odds with negative effects of sourness on liking found in other fruits^34,36^. In hierarchical clustering, liking, strawberry flavor, and sweetness were grouped in one cluster with 100% probability, separate from the sourness cluster with 0% probability (Figure 1).

**Figure 1.**
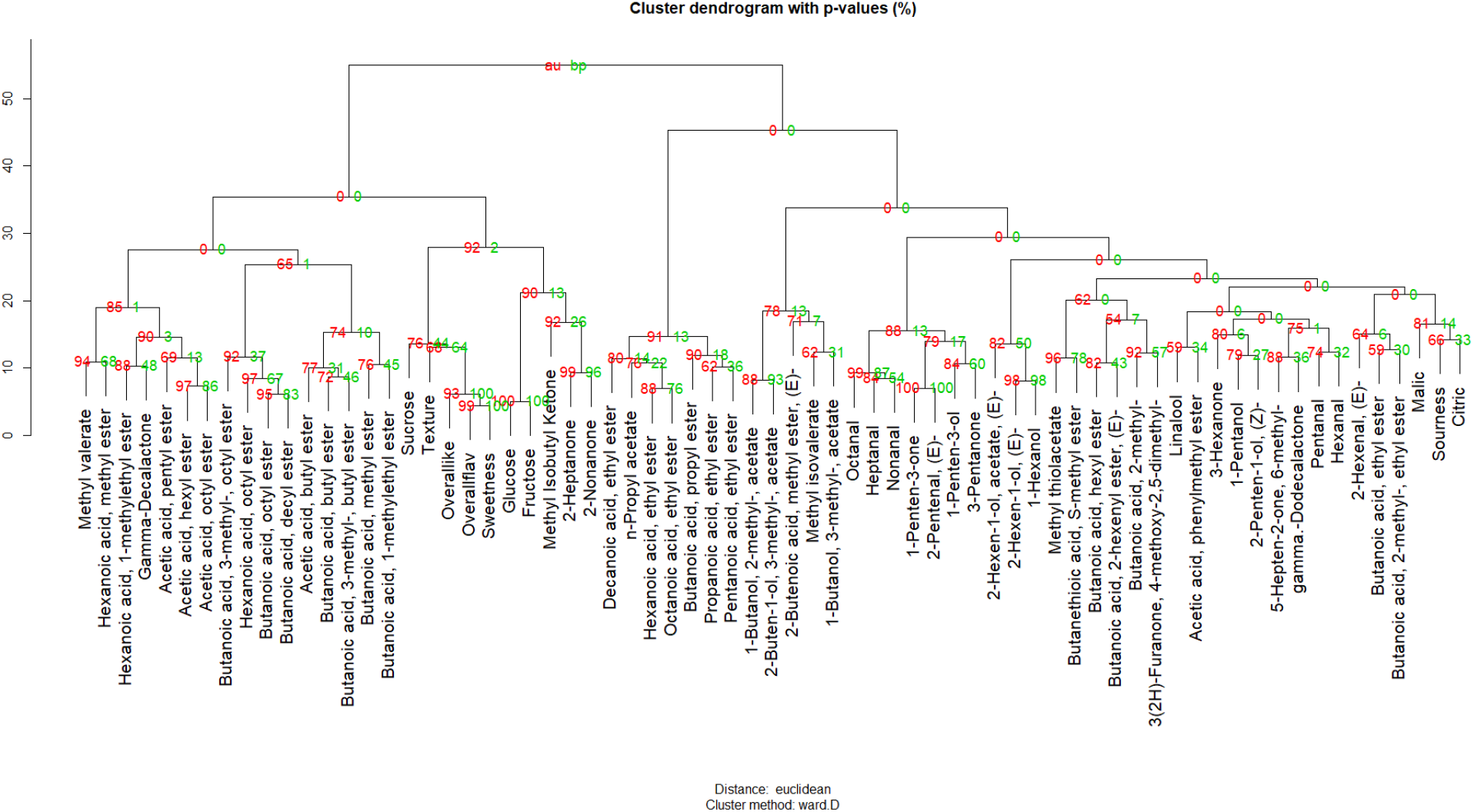
Cluster dendrogram of sensory attributes, volatiles, sugars and acids. AU (Approximately Unbiased) p-values are in red and BP (Bootstrap Probability) p-values are in green.

Due to a diverse collection of strawberry samples from different genetic backgrounds and harvest dates, patterns in perceived sweetness and consumer liking were distinguishable among genotypes and harvest dates. The average liking score was 25.8 across all samples. The top five samples for consumer liking were ‘Festival’ (twice), Sensation® ‘Florida127’, ‘Albion’ and Proprietary 2 (Table S2). However, a large amount of the variability in sensory attributes was driven by temperature. Declines in sweetness and consumer liking over increasing temperature (at different harvest dates) was illustrated with Sensation® ‘Florida127’, ‘Florida Radiance’ and ‘Florida Beauty’ in 2016 and 2017 seasons (Figure 2). The same pattern was also observed for total sugars but not total volatiles (Figure 2). Intersections among lines suggest some level of genotype by environment (G×E) interactions underlying fruit sensory attributes.

**Figure 2.**
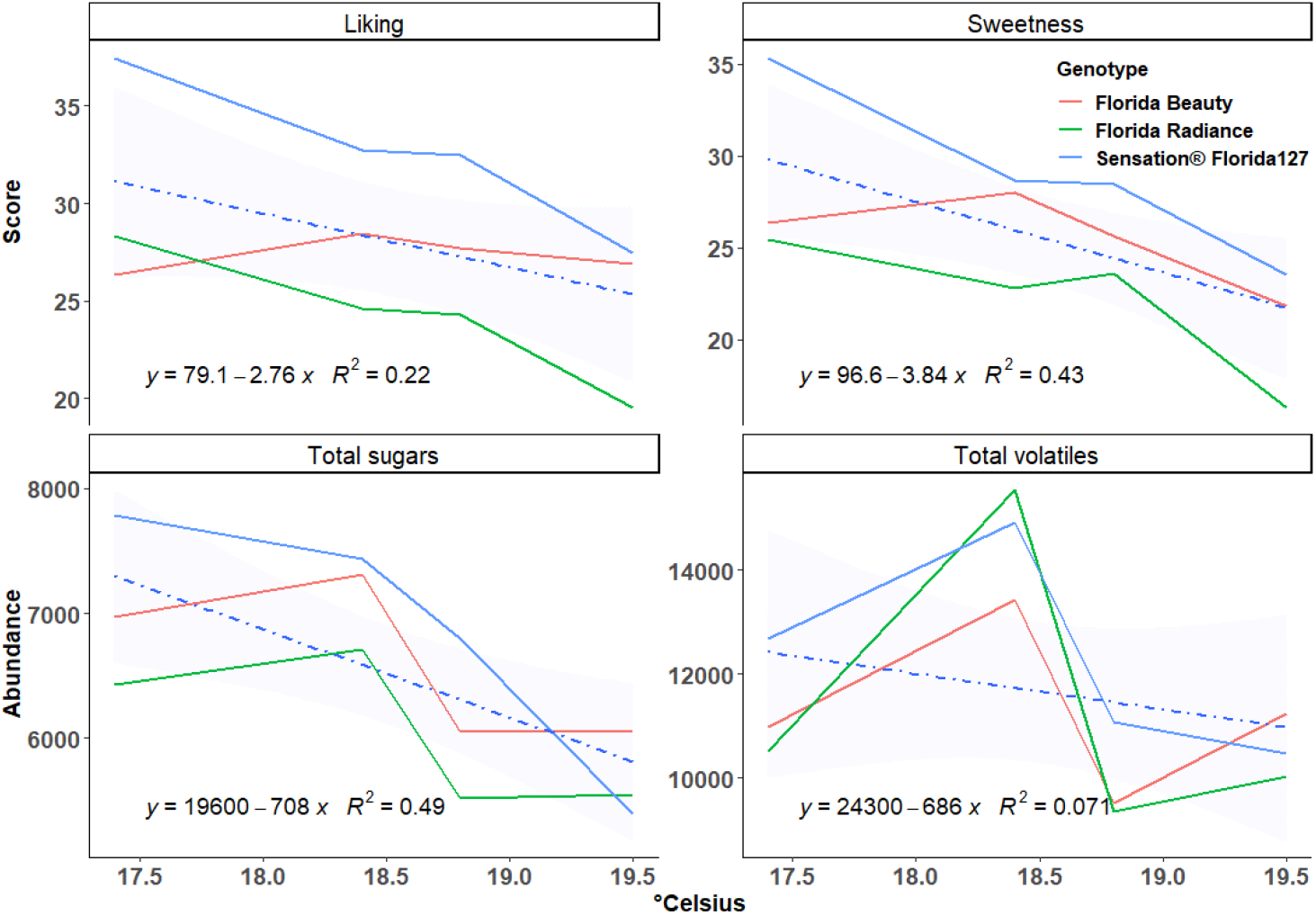
Liking, sweetness, total sugars, and total volatiles of three cultivars as influenced by five-day average soil temperature before harvest. The trend line was modeled with linear regression. Fruits were harvested on 2/22/2016 (five-day average 17.4°C), 3/7/2016 (18.8°C), 2/7/2017 (18.4°C), and 2/14/2017 (19.5°C). Sizable differences among genotypes and among temperatures indicate strong genotype and temperature effects on desired sensory attributes and total sugars.

### Key chemical compounds contributing to sweetness and liking

Chemical analysis of 148 samples yielded 118 compounds including 3 sugars, 2 acids, and 113 volatiles (Table S3). Besides sugars and acids, 59 volatiles were common among all three datasets corresponding to three periods: 2011-2012, 2013-2015 and 2016-2017. Most of the 59 common volatiles were esters (35), followed by aldehydes (7) and ketones (7).

As a first step in identifying the chemical drivers of hedonic experience, correlations among all sensory and chemical attributes were examined. A total of 646 significant correlations (p<0.01) were detected among all variables (Figure 3, Table S4). Glucose (r = 0.63 and r = 0.55) and sucrose (r = 0.62 and r = 0.60) had the highest correlations with both sweetness and liking. In addition to these two sugars, 2-pentenal, (E)- (V12); 1-penten-3-one (V3); heptanal (V37); fructose; butanoic acid, butyl ester (V43); butanoic acid, 3-methyl-, butyl ester (V51); 5-hepten-2-one, 6-methyl- (V42); and γ-dodecalactone (V81) were the compounds with the highest correlations (r > 0.4) to liking. The list of highly correlated volatiles (r > 0.4) with sweetness included those correlated with liking, plus acetic acid, hexyl ester (V46); butanoic acid, octyl ester (V69); acetic acid, butyl ester (V21); and butanoic acid, hexyl ester (V62). The number of volatiles negatively correlated with liking or sweetness was smaller. 2-Hexen-1-ol, acetate, (E)- (v47) exhibited the strongest negative correlations with both liking (r = −0.27) and sweetness (r = −0.21). In a network analysis, significant correlations with liking and sweetness were visualized with edges connecting to sugars; multiple esters; γ-dodecalactone (V81); 5-hepten-2-one, 6-methyl- (V42); 1-penten-3-one (V3); heptanal (V37); 2-pentenal, (E)- (V12) (Figure 4).

**Figure 3.**
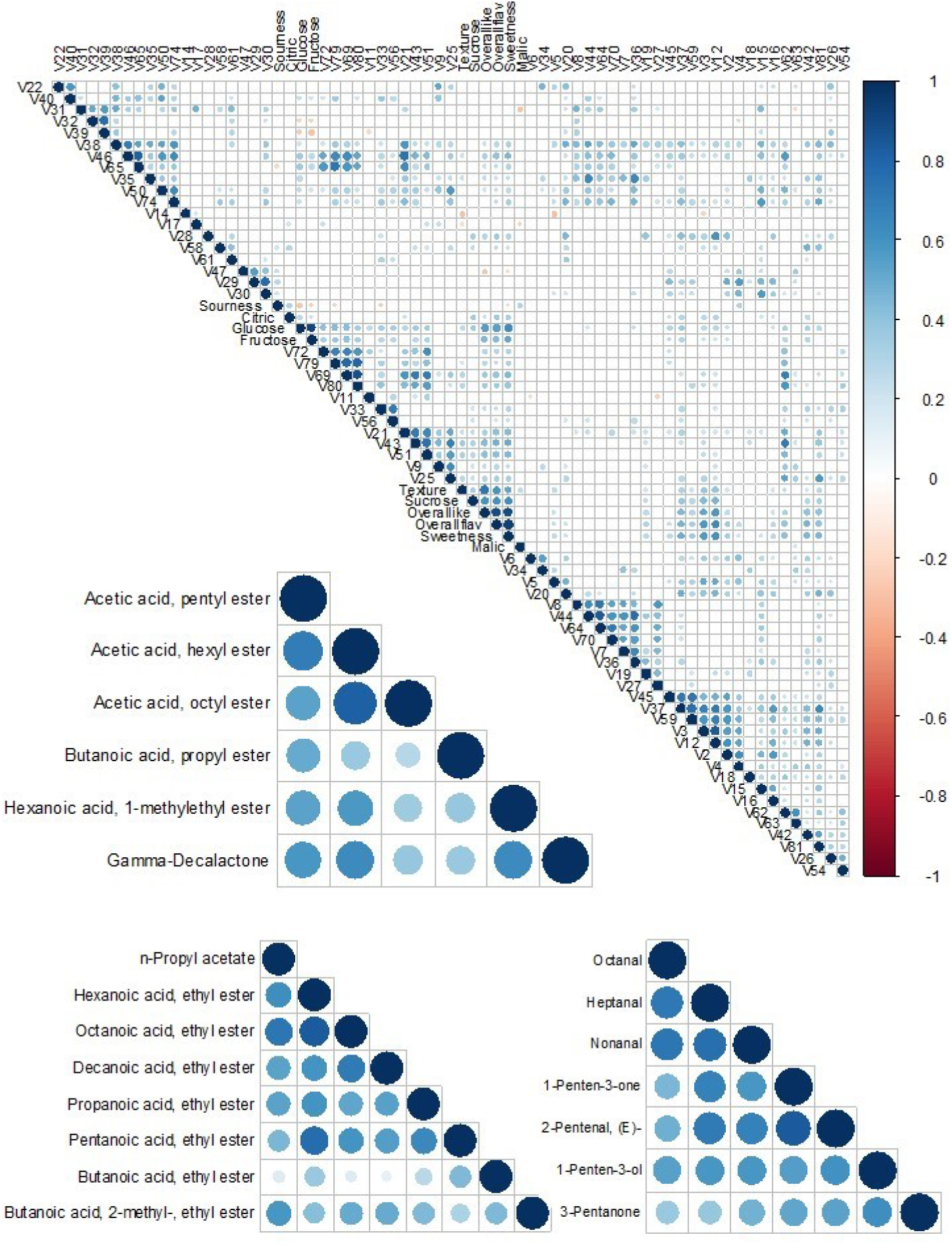
Pairwise correlations of consumer attributes and chemical compounds. Dots represent significant correlations after false discovery rate (FDR) correction. The intensity of color and size of each dot is propotional to the absolute value of the correlation coeffcient (r). Three highly correlated regions of the plot are enlarged.

**Figure 4.**
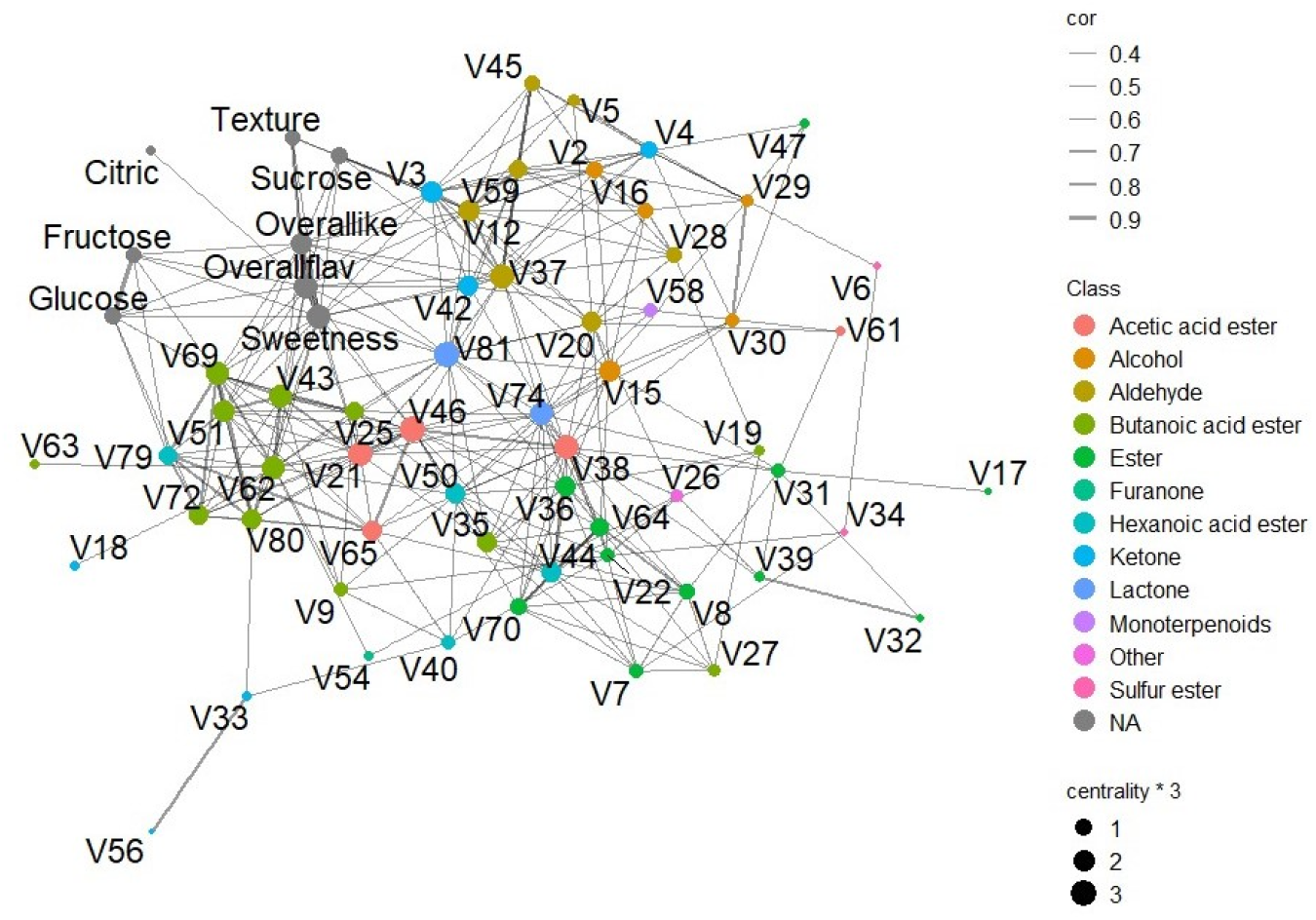
A sensory-chemical network for strawberry. Significant correlations are connected by edges. Sizes of nodes are proportional to centrality scores. Centrality scores were calculated for each node to reveal importance of the volatile in the cluster. Volatiles are colored by chemical class.

Most volatiles significantly correlated with sweetness and liking were corroborated in the PLS models. PLS models were able to explain more than 80 percent of variation for sweetness in each of the data subsets (R^2^=82.93%, 80.96% and 93.37%, respectively). PLS is advantageous when analyzing small samples sizes, and the highest R^2^ was obtained for the 3rd dataset which has the smallest sample size. VIP tests revealed 20 important volatiles (VIP>1) common to the three subsets in addition to glucose, fructose and sucrose (Table 1 and Table S5). These consist of 10 esters, 5 aldehydes, 2 lactones, 2 ketones, and 1 alcohol. Sixteen of these volatiles passed t-tests derived from linear models at Bonferroni corrected p-values < 0.05, indicating that they increased sensory sweetness independent of total sugars. Important esters included seven butanoic acid esters. Among them, butanoic acid, butyl ester (V43) has the highest r of 0.51. Two abundant lactones, γ-dodecalactone (V81) and γ-decalactone (V74) were both selected in the PLS, also showing high correlations with sweetness (r = 0.45 and 0.33). 5-Hepten-2-one, 6-methyl- (V42) with floral aroma was one of the two ketones showing enhancing effect on sweetness with high correlation (r=0.45).

**Table 1.** Variable importance for predicting sweetness and liking. Independent partial least square (PLS) models were fitted for 2011-2012, 2013-2015 and 2016-2017 datasets. Variable importance in projection (VIP) scores measure the importance of each variable in predicting the response variable. A t-test was conducted for each of the chemical compound to determine its significance to model sweetness or liking while controlling for total sugars. ^a^ Seasons. ^b^ P-values were ajusted with bonferroni corrections. ^c^ P-value was not obtained for sugars. ^d^Compound not detected. ^e^Compounds with no VIPs did not pass the threshold of 1 which is the typical cutoff for relevant variable selection.

| Chemical compound | VIPs in sweetness model |  |  |  | VIPs in liking model |  |  |  |
| --- | --- | --- | --- | --- | --- | --- | --- | --- |
|  | 2011-2012<sup>a</sup> | 2013-2015 | 2016-2017 | p-value<sup>b</sup> | 2011-2012 | 2013-2015 | 2016-2017 | p-value |
| Glucose | 1.68 | 2.14 | 1.52 | <sup>c</sup> | 1.19 | 2.18 | 1.65 |  |
| Fructose | 1.60 | 1.62 | 1.79 |  | 1.00 | 1.65 | 1.80 |  |
| Sucrose | 2.17 | 2.10 | 0.27 |  | 2.19 | 2.32 | 0.20 |  |
| 1-Penten-3-one | 1.76 | 1.26 | 1.34 | 0.0000 | 1.87 | 1.31 | 1.23 | 0.0000 |
| 2-Pentenal, (E)- | 1.52 | 1.53 | 1.76 | 0.0000 | 1.83 | 1.48 | 1.71 | 0.0000 |
| 2-Penten-1-ol, (Z)- | 0.19 | 1.12 | 2.00 | 1.0000 | 0.33 | 1.02 | 1.94 | 1.0000 |
| Heptanal | 1.36 | 1.26 | 1.60 | 0.0000 | 1.43 | 1.13 | 1.38 | 0.0000 |
| 5-Hepten-2-one, 6-methyl- | 1.16 | 1.42 | 0.49 | 0.0000 | 1.31 | 1.20 | 0.41 | 0.0000 |
| Butanoic acid, butyl ester | 1.38 | 1.48 | 0.88 | 0.0013 | 1.17 | 1.48 | 1.03 | 0.3781 |
| Acetic acid, hexyl ester | 1.39 | 1.24 | 0.39 | 0.0013 | 1.17 | 1.12 | 0.52 | 1.0000 |
| Butanoic acid, 3-methyl-, butyl ester | 1.23 | 1.33 | 0.77 | 0.0000 | 1.05 | 1.42 | 1.02 | 0.0033 |
| Butanoic acid, pentyl ester | 1.28 | 1.18 | 0.00 | 0.0000 | 1.28 | 1.12 |  | 0.0000 |
| Nonanal | 1.16 | 0.81 | 1.36 | 0.0000 | 1.33 | 0.77 | 1.30 | 0.0013 |
| Butanoic acid, hexyl ester | 1.50 | 0.90 | 1.71 | 0.0467 | 1.27 | 0.87 | 1.70 | 1.0000 |
| Butanoic acid, octyl ester | 1.01 | 1.30 | 1.30 | 0.0007 | 0.79 | 1.32 | 1.40 | 0.2430 |
| Butanoic acid, decyl ester | 1.06 | 1.03 | 1.23 | 0.0000 | 1.01 | 1.07 | 1.27 | 0.0005 |
| γ-Dodecalactone | 1.51 | 1.25 | 0.74 | 0.0000 | 1.54 | 1.21 | 0.84 | 0.0000 |
| Hexanoic acid, butyl ester | <sup>d</sup> | 1.34 | 1.44 | 0.0002 |  | 1.26 | 1.48 | 1.0000 |
| γ-Decalactone | 1.08 | 1.12 | 0.69 | 0.2637 | <sup>e</sup> | - | - | - |
| Acetic acid, butyl ester | 1.16 | 1.34 | 0.44 | 0.0371 | - | - | - | - |
| 2-Hexenal, (E)- | 1.17 | 0.52 | 1.61 | 0.0000 | - | - | - | - |
| Benzaldehyde |  | 1.01 | 1.99 | 1.0000 | - | - | - | - |
| Butanoic acid, phenylmethyl ester |  | 1.03 | 1.04 | 1.0000 | - | - | - | - |
| Butanoic acid, propyl ester | - | - | - | - | 1.18 | 1.09 | 0.86 | 1.0000 |
| Methyl Isobutyl Ketone | - | - | - | - | 0.79 | 1.07 | 1.47 | 1.0000 |
| Butanoic acid, 3-methyl-, octyl ester | - | - | - | - | 0.91 | 1.21 | 1.05 | 1.0000 |

A slightly lower portion of variation in response was explained in the liking models (77.56%, 70.49% and 92.41%, respectively). Sucrose was the most important compound to consumer liking in the first two datasets. Fructose and glucose had lesser but important influence on liking in all three datasets. Important volatiles obtained for consumer liking included 10 esters, 3 aldehydes, 1 lactone, 3 ketones, and 1 alcohol. Eleven of these compounds were significant for the t-test (Table 1 and Table S5), providing evidence that they had positive impact on consumer liking that was independent of total sugars. Butanoic acid, ethyl ester (V19), known to be one of the most abundant and important contributors to strawberry aroma, showed significant positive correlations (r =0.31) with liking but did not pass the PLS threshold for two of the three datasets. Other abundant esters like butanoic acid, methyl ester (V9); hexanoic acid, methyl ester (V40) and hexanoic acid, ethyl ester (V44) all showed positive but low to moderate correlations (r=0.29, 0.17, and 0.13, respectively) with liking and did not reach PLS thresholds. Fifteen compounds were identified via PLS as important drivers of both sweetness and liking models, reflecting the high correlation between sweetness and liking (Table 1).

### Predicting sweetness and liking with sugars, acids, and volatiles

Since sugars and acids are known to be important factors influencing sweetness perception and consumer liking, we first built a model with sugars and acids. This model produced an R^2^ of 0.41 in testing CV (independent test dataset) for sweetness, indicating that 41% of variation in sweetness could be accounted for by sugars and acids. To further investigate the influences of volatiles on sweetness and liking, we included them as variables and tested prediction accuracies with a broad spectrum of algorithms (Figure 5a), including decision-tree based Random Forest (RF), bayes GLM, GLM, GLMBOOST, and LASSO. With the inclusion of volatiles, LASSO and GLMBOOST models explained the most variation for sweetness in testing CV (R^2^ = 0.65 and 0.69), outperforming the model with sugars and acids by 28% for the test dataset. In addition, LASSO and GLMBOOST generated the lowest standard deviation of R^2^ among all models (Figure 5a). Similar results were produced in prediction of liking but with a slightly lower total variance explained. LASSO model accounted for 55% in testing CV, compared to only 30% in models with sugars and acids alone (Figure 5b). These results provide further strong evidence of volatile compound enhancement of strawberry sweetness perception and consumer liking independently of sugars.

**Figure 5.**
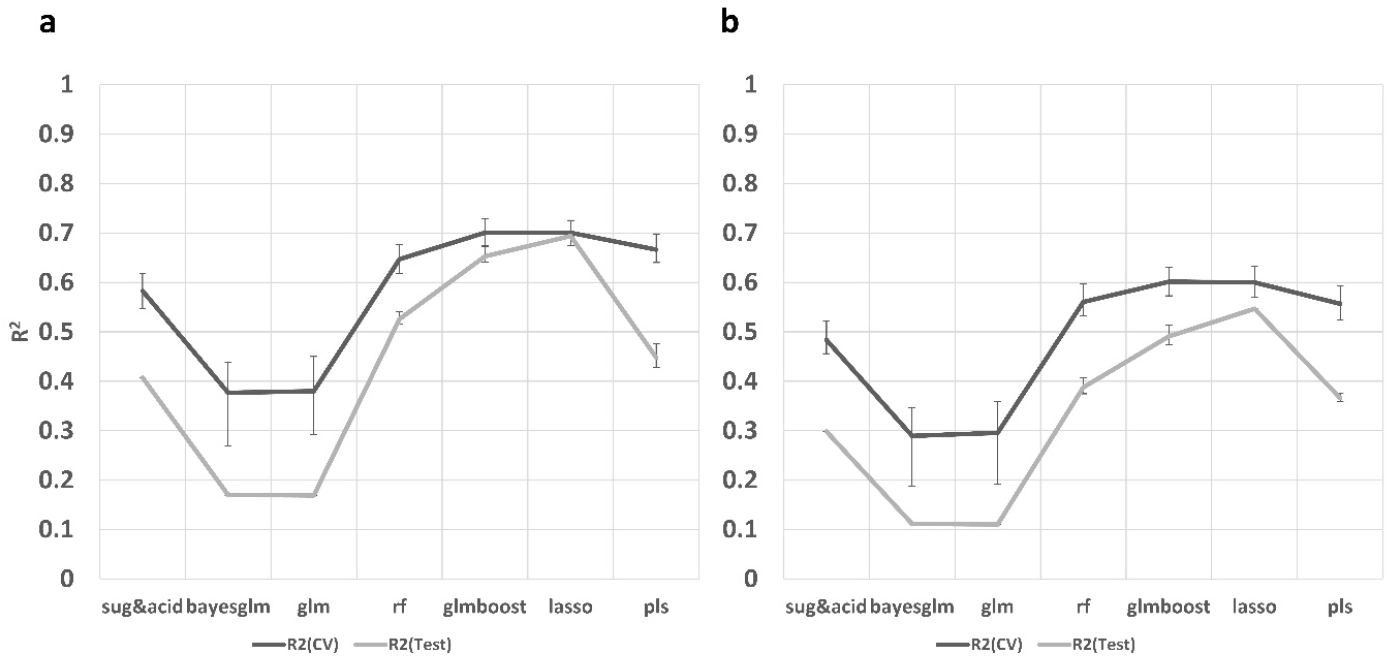
Histograms of performance of six machine learning algorithms using sugars, acids, and volatiles to predict sweetness intensity (**a**) and consumer liking (**b**) compared to a model with sugars and acids (leftmost in both figures). Black lines represent the average R^2^ of nested cross validation while grey lines show the average R^2^ when using a test dataset. Error bars represent standard deviations based on 100 iterations.

### Chemical networks in strawberry fruit

Broad variations in quantity of chemical compounds were measured for the 148 strawberry samples. For example, the most abundant ester (butanoic acid, methyl ester; V9) ranged from 92.0 ng^1^ 100gFW^-1^ hr^-1^ in ‘Radiance’ in 2017 to 7359.6 ng^1^ 100gFW^-1^ hr^-1^ in a proprietary sample from 2011, an approximately 70-fold difference. Qualitative differences were also observed. For example, methyl anthranilate (V68) was only detectable in 3 samples and 2-nonanone (V56) was detected in 99 of 148 samples (Table S3).

Collinearity among chemicals may reflect metabolic pathways, helping to narrow the number of genetic targets for flavor improvement. Fructose and glucose exhibited a very high correlation (r = 0.91). The highest positive correlations among volatiles were found between butanoic acid, octyl ester (V69) and butanoic acid, decyl ester (V80) (r = 0.87), followed by hexanoic acid, ethyl ester(V44) and octanoic acid, ethyl ester (V64) (r = 0.83); 1-penten-3-one (V3) and 2-pentenal, (E)- (V12) (r = 0.83); and acetic acid, hexyl ester (V46) and acetic acid, octyl ester (V65) (r = 0.82). Other correlations are detailed in Figure 3 and Table S4.

Group correlations and clusters among volatiles were better illustrated with hierarchical clustering and network analyses. Fatty acid esters derived from the same alcohol were grouped together in several clusters (Figure 1). For example, acetic acid, butyl ester (V21); butanoic acid, butyl ester (V43); and butanoic acid, 3-methyl-, butyl ester (V51) were grouped with 81% confidence. Butanoic acid, 3-methyl-, octyl ester (V72); hexanoic acid, octyl ester (V79); butanoic acid, octyl ester (V69); and butanoic acid, decyl ester (V80) were clustered with 93% confidence. Other fatty acid ester clusters contained esters derived from the same fatty acid. For instance, acetic acid, pentyl ester (V38); acetic acid, hexyl ester (V46); and acetic acid, octyl ester (V65) were grouped with 62% confidence. A sulfur ester group was remote from the main ester group with a strong bond (96% confidence) between methyl thioacetate (V6) and butanethioic acid, S-methyl ester (V34). Besides esters, we observed an aldehyde group including octanal (V45), heptanal (V37) and nonanal (V59) with 99% confidence.

In the chemical networks, sixty-five nodes were connected by 299 significant edges (Figure 4). A distinct butanoic acid ester cluster consisting of 8 esters including butanoic acid, hexyl ester (V62) and butanoic acid, octyl ester (V69). A second ester cluster centered on hexanoic acid, ethyl ester (V44) included multiple ethyl esters, such as octanoic acid, ethyl ester (V64), pentanoic acid, ethyl ester (V36), and propanoic acid, ethyl ester (V7). Aldehydes had their own cluster around heptanol (V37), surrounded by 2-pentenal, (E)- (V12); nonanal (V59); hexanal (V20); pentanal (V5); octanal (V45) and 2-hexenal, (E)- (V28).

### Genetic and chemical diversity among tested varieties

In order to explore the genetic diversity of the genotypes tested, we obtained SNP data via the Axiom® IStraw35 array^37^ for 26 samples. The genotyped panel comprised 20 UF genotypes, 4 UC-Davis genotypes, 1 proprietary genotype and 1 genotype of European origin. The PCA individual plot showed a clear separation between UC and UF genotypes on PC2 (Figure 6a). On the other hand, PC1 explained the most variation within UF genotypes. Combining PC1 and PC2 explained 22.1 percent of variation. In a complementary analysis we conducted PCA with sugars, acids and 59 common volatiles (Figure 6b). No visible separation for genotypes from different origins was observed in the individual plot. Combining PC1 and PC2 explained 29 percent of variation.

**Figure 6.**
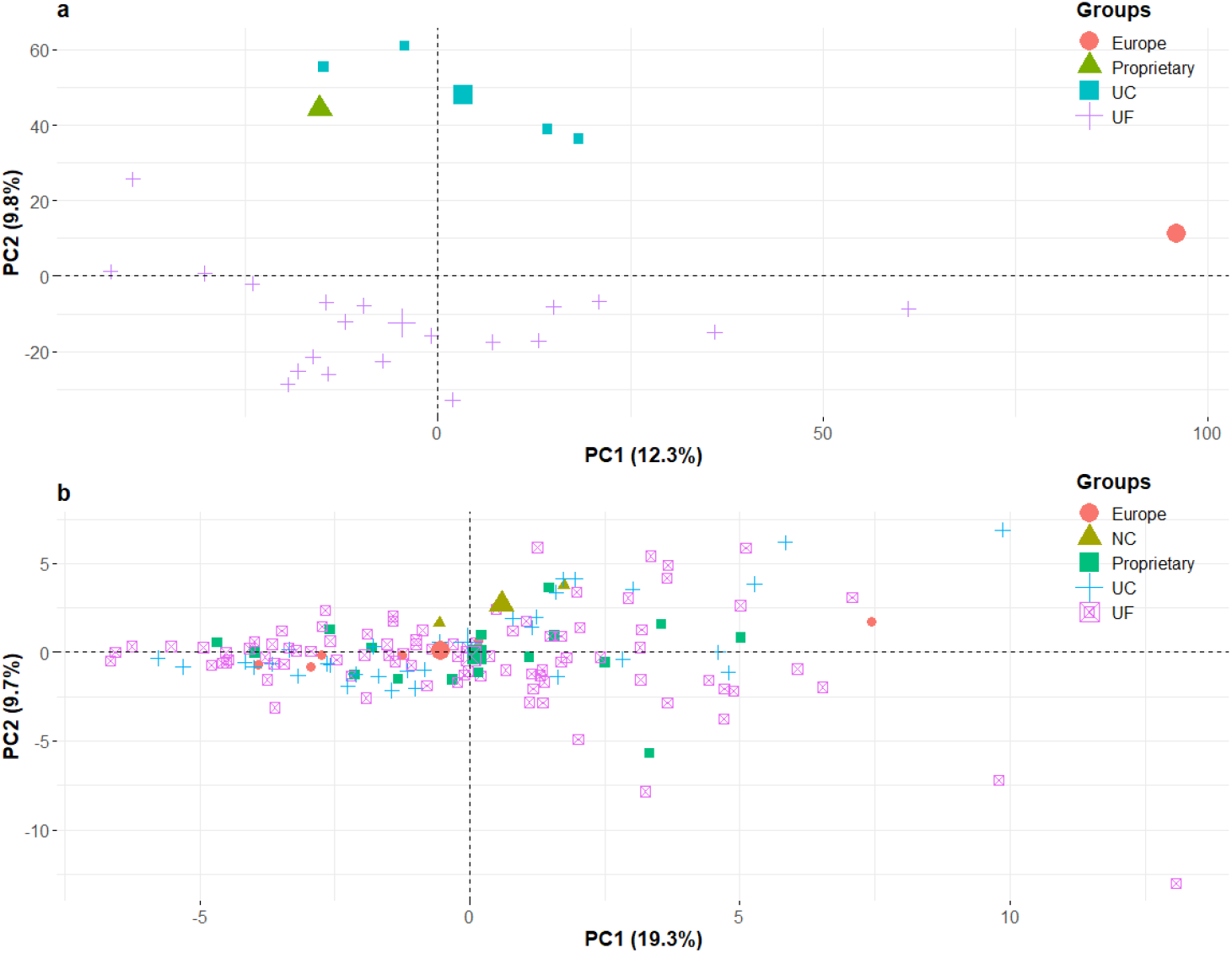
(**a**) Principal component analysis (PCA) plot of genotypes using Axiom® IStraw35 38000 single nucleotide polymorphisms (SNPs). Genotypes from different origins are grouped by color. Percentage of total variation explained by each component is shown in parentheses. (**b**) PCA plot of genotypes using volatiles, sugars, and acids.

### Two major QTLs for production of esters

A chemical network analysis suggested the possibility of common genetic control for multiple esters (Figure 4). To investigate this possibility, QTLs for ester production were mapped across eight experimental crosses. Medium-chain esters including acetic acid, decyl ester; acetic acid, hexyl ester; acetic acid, octyl ester; acetic acid, nonyl ester; butanoic acid, butyl ester; and butanoic acid, octyl ester shared a single QTL on linkage group 6A (Figure S1). Single-marker analyses revealed that a shared maker (AX-123358920) explained 7.7% to 30.3% of the variation in ester abundances. Another QTL on the same linkage group but at a distinct location was discovered for both butanoic acid, ethyl ester and hexanoic acid, ethyl ester (Figure 7). The single shared marker (AX-166508266) explained 15.0% of the observed variance in hexanoic acid, ethyl ester abundance (p = 1.5e-5) and 15.6% of butanoic acid, ethyl ester abundance (p = 1.6e-5).

**Figure 7.**
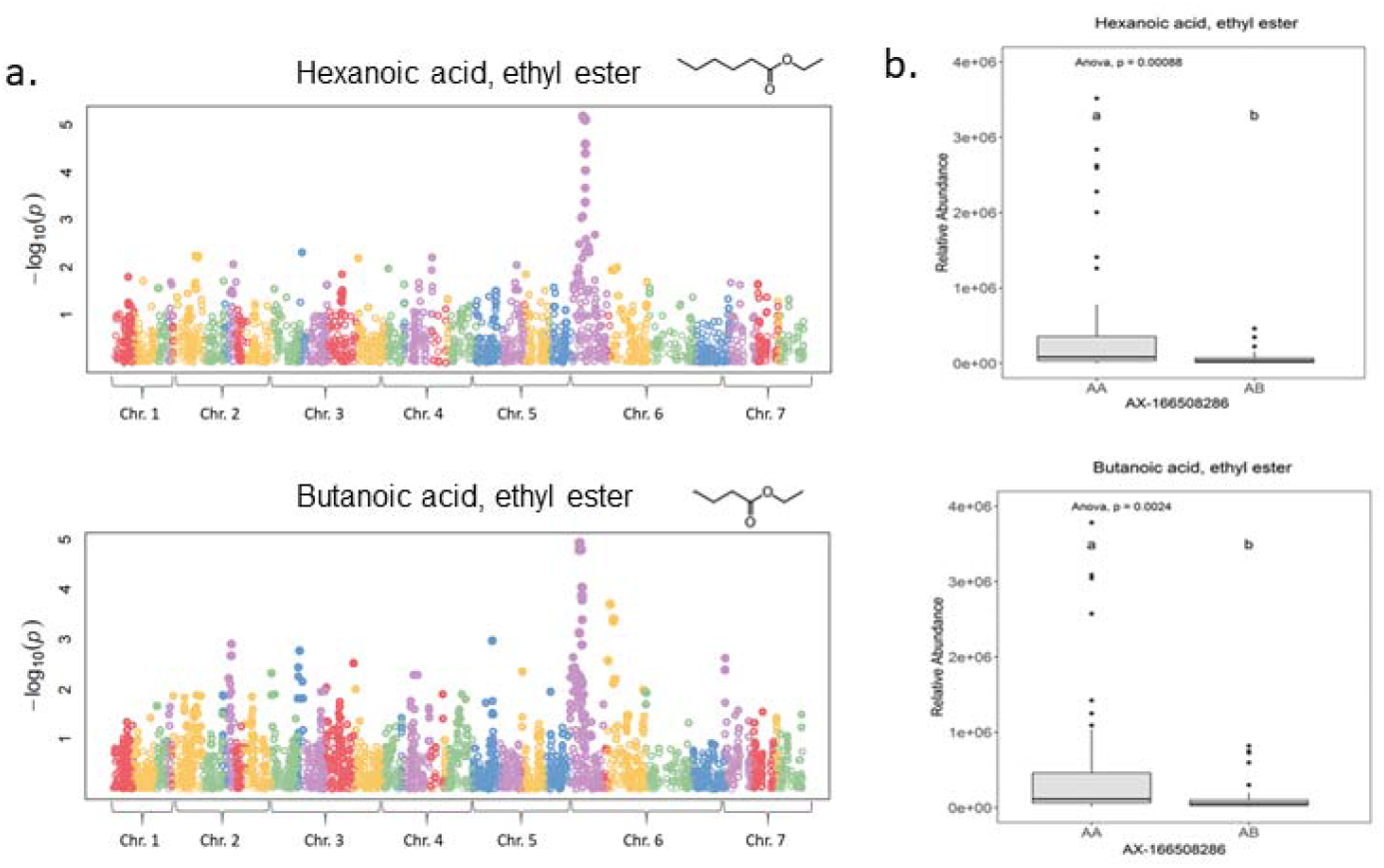
(a) Manhattan plots for hexanoic acid, ethyl ester and butanoic acid ethyl ester. A shared QTL was observed on linkage group 6A. (b) A single shared marker explained a large portion of variation in the corresponding ester abundance. Different letters indicate significant differences at p = 0.05.

## Discussion

A large-scale sensory and chemical analysis of 148 fresh strawberry samples was conducted over seven years for the purposes of examining the sensory and chemical drivers of consumer preference. Our study largely expands the previous work^18^ in terms of sample size, and provided new insights towards volatiles contributing to liking, chemical network and clustering, and volatile genome-wide association study, most of which were not generalizable if using a small sample collection. In agreement with studies of tomato and blueberry and the previous strawberry work, perceived sweetness (r = 0.714*) was the major determinant for liking in strawberry^15,18,34^. A previous study using a glucose solution found that sweetness and hedonics followed a biphasic relationship in which the apex of liking was reached at glucose concentration of 500 mM^38^. In the present study the highest recorded total sugars concentration was 10056.2 mg^1^100gFW^-1^, roughly estimated to approximately 10 mM glucose solution, much lower than the equivalent optimal concentration. This was consistent with a low degree of satisfaction in sweetness in the sensory panel, where only 8.2% of strawberry samples reached the panelist’s reported ideal sweetness (data not shown). This means that 91.8% of samples the sensory panels failed to achieve the panelist’s ideal sweetness for a strawberry. This discrepancy between ideal sweetness and sample sweetness dramatically illustrates the need for breeding to increase sweetness in strawberries. One strategy to enhance perceived sweetness is to increase sugar concentration. The major sugars in strawberry fruit are glucose, fructose and sucrose^39^. All three sugars exhibited high correlations with sweetness (Table1 and Figure 3) and were the top drivers of liking. Nevertheless, breeding strawberries for higher sugar content may result in negative impacts on other critical traits. A trade-off between yield and soluble solids content (SSC) was found in UF strawberry breeding germplasm at the additive genetic level, demonstrating that genetic gains for SSC would likely result in yield losses^40^. In peach, higher SSC is associated with a longer fruit development period and reduced fruit weight^41^. In contrast, volatiles typically occur at 10^3 to 10^6 times lower concentration (uM to nM range) than sugars. Increasing sweetness-enhancing volatile concentrations even a small amount has the potential to increase the perception of sweetness and overall liking at a lower carbon cost to industry-demanded agronomic traits. Thus, non-sugar sources of sweetness would be highly advantageous for improving both flavor and agronomic traits simultaneously.

When fruits are chewed or swallowed, volatiles flow into the retronasal tract. Signals from retronasal olfaction interact with signals from the tongue in the brain, imparting the sensation of taste^15^. Evidence of integration of flavor and taste was adduced by a faster response time when drinking a flavored sugar solution^42^. While the effects of volatiles on flavor is widely known, the enhancement of sweetness by specific volatiles in sugar solutions or fruit puree is also a known phenomenon^43^. When accompanied by strawberry odor, perceived sweetness increases at a range of sugar concentrations in water solution^44^. In the present study, we demonstrate this phenomenon via consumer testing of actual strawberry fruit.

Strawberry volatile compounds are highly diverse and variable. A recent study consolidating information from 25 strawberry volatile studies from 1991 to 2016 found no consensus on the detected volatiles^5^. The most frequently identified volatile compounds have been distributed among the esters, acids, lactones, aldehydes, furans, alcohols, ketones, and terpenoids. In our chemical analyses, we captured 113 volatiles with 59 common to all years (Table S3). In our previous study comprising 54 of these samples, six volatiles (1-penten-3-one (V3); γ-dodecalactone (V81); butanoic acid, pentyl ester (V53); butanoic acid, hexyl ester (V62); acetic acid, hexyl ester (V46); and butanoic acid, 1-methylbutyl ester (V48)) were found significantly correlated with sweetness independent of sugars^18^. In the present study, five of these six volatiles except for butanoic acid, 1-methylbutyl ester were validated in additional datasets and fourteen additional sweetness-enhancing volatiles and eighteen liking-enhancing volatiles were reported. Among the nineteen sweetness enhancers, ten are esters with nine of the ten having a carbon number more than eight. To our surprise, the most abundant esters butanoic acid, methyl ester (V9) and butanoic acid, ethyl ester (V19) did not reach the VIP threshold for some datasets, even though significant correlations were found between butanoic acid, ethyl ester and liking and sweetness.

Strawberry varieties with high concentrations of lactones, which impart peach-like odor, have been implicated as preferable in previous consumer tests^45^. We show evidence that γ-decalactone (V74) and γ-dodecalactone (V81) are sweetness-enhancing. Nevertheless, only γ-dodecalactone was important for liking. Distributions of the two lactones were also disparate, with γ-decalactone following a bimodal distribution in our study, consistent with its presence being controlled by a single gene^19^. On the other hand, γ-dodecalactone followed a normal distribution suggestive of polygenic control. Three correlated volatile compounds, 1-penten-3-one (V3); 2-pentenal, (E)- (V12); and 2-penten-1-ol, (Z)- (V16) were highly associated with sweetness and liking. Although none of them elicit sweet aroma according to the literature^46^, their interactions with sugars or other volatiles at the neurological level may lead to increased sweetness perception and hedonic experience. Interestingly, 1-penten-3-one and 2-pentenal, (E)-positively impacts liking in tomato^16^, indicating that common contributors to liking may exist across a wide diversity of fruits. Two volatiles impacting liking in this study, nonanal (V59) and 5-hepten-2-one, 6-methyl- (V42) impart flowery notes to the strawberry, which are important building blocks in the diversity of strawberry aroma. Carotenoids including 5-hepten-2-one, 6- methyl- were also positively correlated with consumer acceptability in tomato^47^. Due to the isolation and the detection techniques employed in this study, furaneol was not quantified. Future experiments investigating the influence of furaneol on strawberry will need to harness other isolation approaches. In tomato, Buttery *et al.*^48^ demonstrated the isolation of and detection furaneol with anhydrous sodium sulfate coupled with dynamic headspace sampling.

Fruit chemical data have great potential for predicting sensory responses and carry the advantage of accuracy and objectivity as targets for genetic improvement. Univariate models with soluble solid content or titratable acidity have often provided poor prediction of liking or sensory attributes^49^. Including additional sensory contributors like volatiles as variables can increase model prediction accuracy. However, a mathematical challenge of adding volatile data to prediction models is the depletion of degrees of freedom with high dimensional data. A few studies have addressed this problem by adopting different statistical models like multivariate models, PLS models, principle component regression models, and artificial neural networks^35,49,50^. In these studies, machine learning models always resulted in higher explained variance of sensory attributes than simple models with sugars only. However, these studies have limited applicability, suffering from small sample size or lack of cross-validation^50,51^. In this study, 90% samples were used to train the model and 10% was exclusively used for testing. Comparisons of multiple prediction methods revealed a 28% increase in variation explained for sweetness using LASSO model compared to linear model with sugars and acids, as well as 25% in liking. GLMBOOST and LASSO models exhibited better performance than the traditional PLS model in all quality measurements. On the other hand, the large increasement of R-square value when incorporating volatile data corroborates the importance of volatiles to enhance sweetness and consumer liking.

Strawberry quality is strongly influenced by both genetic and environmental factors^5,25,52^. In our analyses, genetic variation affected presence and absence of volatiles as well as their abundance (Table S3). In addition, temperature fluctuated dramatically during the evaluated seasons, from periods of nearly freezing weather in December and January to an average over 26 °C in March and April, far above the optimal temperature for fruit quality^52^. Thirteen compounds including citric acid and total sugars were significantly affected by harvest date within season (data not shown). Similarly, multiple other studies found declines of total sugars or SSC during the course of strawberry growing seasons^25,53^. Although a seasonal decline was observed for total volatile abundance (Figure 2), it was not statistically significant, and not all volatiles were negatively affected by temperature. In blueberry, fructose and glucose abundance exhibited significant G×E or environmental effects, in contrast to many of the volatiles^34^. Perhaps volatiles may hold part of the key for stabilizing sensory qualities in the face of a changing environment and may attenuate the impact of declines in total sugars with increasing temperature. Network and clustering analyses of metabolites may provide evidence for common biosynthetic pathways or regulation mechanism to help narrow targets for genetic improvement. Here we used pair-wise correlations and hierarchical clustering approach to explore connections among volatiles. Fatty acid esters were clustered based on either alcohol substrate or acyl-CoA substrate. Butanoic acid esters were clustered by both approaches with high confidence (Figure 1 and 4). Ethyl esters, such as octanoic acid, ethyl ester (V64); pentanoic acid, ethyl ester (V36); propanoic acid, ethyl ester(V7); and hexanoic acid, ethyl ester (V44) were highly correlated and clustered in the volatile network. Common genetic regulation for esters was further corroborated with the discovery of major QTLs shared among multiple medium-chain esters and the two most abundant ethyl esters. In a European strawberry mapping population, possibly the same QTL on linkage group 6-1 influenced the productions of multiple medium-chain esters^54^, which supports broad applications for maker assisted selection at this locus to improve sensory sweetness and flavor. The potential candidate gene underlying the ester QTL hotspot might be an alcohol acyltransferase (AAT). AATs catalyzing the final steps of ester synthesis have diverse functional paralogs in octoploid strawberry^55,56^. Some important esters for liking derived from medium-chain aliphatic alcohols match the primary products of SAAT^57^. Another important volatile class, aldehydes, synthesized via decarboxylases or hydroperoxide lyase^58,59^, has not been well-characterized in strawberry. Our network analysis also predicts a common enzyme catalyzing a range of aldehydes. In wild relative *Fragaria vesca,* a volatile metabolomics study using near-isogenic lines also showed ester and aldehyde clusters^60^. Similar coordinated metabolic networks were demonstrated in peach, tomato, apple, and melon^15,61–63^. Together these analyses reveal concerted regulation within and between volatile classes in both climacteric and non-climacteric fruits.

While consumers desire many traits in fresh strawberries, including appearance and health attributes, flavor ranks at the top^2^. Flavor is a difficult trait to improve due its chemical complexity, strong environmental effects and negative relationships with grower-demanded agronomic traits. It is also not trivial to measure. Quantifying sensory attributes for every genotype in the breeding pool is not practical, nor is comprehensive volatile quantification. In this study we have identified a reduced set of chemical targets for consumer liking and sweetness improvement in strawberry. High-throughput and cost-effective DNA markers can be easily developed based on the significant makers identified from our association analysis and implemented in diverse strawberry breeding programs. We must also consider that this study intentionally samples the chemical diversity of commercial germplasm that is readily deployed in agriculture. Yet distinct flavors in the wild relatives of cultivated strawberry could be introgressed into the cultivated germplasm pool, expanding the flavor toolbox of modern strawberry in the long-term. Narrowing the non-sugar targets for sweetness-enhancement in strawberry is an important step toward a worthy goal: increased consumer satisfaction for fresh strawberries.

## Materials and Methods

### Ethics statement

Consumer panels were conducted at the Food Science and Human Nutrition Department at the University of Florida (UF) in Gainesville, FL. The UF Institutional Review Board 2 (IRB2) chaired by Ira S. Fischler approved the protocol and written consent form (case #2003-U-0491) that sensory panel participants were required to complete.

### Plant material and fruit production

From 2010-2017, 148 strawberry (*Fragaria ×ananassa*) samples from 48 cultivars and breeding lines (Table S6) were harvested from strawberry research plots of the University of Florida (UF) Gulf Coast Research and Education Center (GCREC) (Balm, FL) or the Florida Strawberry Growers Association (FSGA) headquarters in Dover, FL. All fruiting field trials were managed according to commercial standards for Florida annual plasticulture^64^. Data from 54 samples harvested during 2010-2012 were analyzed and published in a previous paper^18^. These samples represented major commercial varieties grown in California, Florida and Europe. Ninety-four additional samples were collected during 2013-2017, representing a wider range of UF germplasm including varieties and advanced breeding lines. Thirty-three genotypes were harvested more than one time across the seven years. The combined 7-year dataset captured variation produced by genetics, harvest dates and seasons (Table S6).

At each harvest date, fully ripe fruit with at least 90% red surface from three to five genotypes were harvested from the field in the early morning, transported to the Food Science sensory lab in Gainesville, FL, and stored at 4°C in the dark overnight. Each sample consisting of at least 200 fruits was collected from a set of 500 plants. Seven or more fruits, each collected from a separate plant were randomly chosen for chemical quantification. The rest were used for sensory evaluation during the next morning. Temperature data were obtained from the Balm, FL station of the Florida Automated Weather Network located directly adjacent to the plots from which strawberries were sampled. Soil temperature was recorded at a depth of −10 cm. Temperature data were used to regress liking, sweetness, total sugar and total volatiles from three cultivars during the 2016 and 2017 seasons.

### Sensory evaluation

In each session, an average of 100 consumers were divided into individual booths and utilized specialized sensory software to rate the samples (CompuSense® 5 Sensory Analyses Software, CompuSense, Guelph, Canada). One or two whole strawberries from each of three to five randomly coded and ordered genotypes were served per individual/session. All strawberries were served at room temperature.

Consumers were asked to rate hedonic attributes (consumer liking and texture liking) on the Global Hedonic Intensity Scale (GHIS), which was anchored with the most intense liking ever experienced at the top (+100) and the most intense disliking ever experienced at the bottom (−100). Intensity of individual sensory attributes (sweetness, sourness and strawberry flavor) were scored on the Global Sensory Intensity Scale (GSIS) which is anchored with the most intense sensation at the top (+100) and no sensation at the bottom (0)^65^.

### Volatile detection and quantification

At least seven fruits were removed from 4°C storage and pooled in a blender prior to splitting into three 15 g replicates for immediate capture of volatile emissions. The remainder was frozen in liquid nitrogen and stored at −80°C for subsequent quantification of sugars and acids. A two-hour collection in a dynamic headspace volatile collection system^66^ allowed for concentration of emitted volatiles on HaySep 80–100 porous polymer adsorbent (Hayes Separations Inc., Bandera, TX, USA). Elution from the polymer was described by Schmelz^67^. Quantification of volatiles in an elution was performed on an Agilent 7890A Series gas chromatograph (GC) (carrier gas; He at 3.99 ml min-1; splitless injector, temperature 220°C, injection volume 2 μl) equipped with a DB-5 column ((5%-phenyl)-methylpolysiloxane, 30 m length ×250 μm i.d. × 1 μm film thickness; Agilent Technologies, Santa Clara, CA, USA). Oven temperature was programmed from 40°C (0.5 min hold) at 5°C min^-1^ to 250°C (4 min hold). Signals were captured with a flame ionization detector (FID) at 280°C. Peaks from FID signal were integrated manually with Chemstation B.04.01 software (Agilent Technologies, Santa Clara, CA). Volatile emissions (ng^1^ 100gFW^-1^ h^-1^) were calculated based on individual peak area relative to sample elution standard peak area. GC-mass spectrometry (MS) analyses of elutions were performed on an Agilent 6890N GC in tandem with an Agilent 5975 MS (Agilent Technologies, Santa Clara, CA, USA), and retention times were compared with authentic standards (Sigma Aldrich, St Louis, MO, USA) for volatile identification^68^.

### Sugars and acids quantification

Titratable acidity, pH, and soluble solids content were averaged from four replicates of the supernatant of centrifuged and thawed homogenates^69^. An appropriate dilution of the supernatant from a separate homogenate (centrifugation of 1.5 ml at 16,000 x g for 20 min) was analyzed using biochemical kits (per manufacturer’s instructions) for quantification of citric acid, L-malic acid, D-glucose, D-fructose, and sucrose (CAT# 10-139-076-035, CAT# 10-139-068-035, and CAT# 10-716-260-035; R-Biopharm, Darmstadt, Germany) with absorbance measured at 365 nm on an Epoch Microplate Spectrophotometer (BioTek, Winooksi, VT, USA). Metabolite average concentration (mg^1^ 100gFW^-1^) was determined from two to six technical replicates per pooled sample. Derived sucrose concentrations via D-glucose and D-fructose were mathematically pooled.

### Statistical analyses and model parameters

For the purpose of comparing averaged sensory features between samples assessed by different panelists, a mixed linear model was constructed with panelist as a random effect. Least square means (Table S2) of the sensory features for each sample were obtained using SAS software (Version 9.2.; SAS Institute, Cary, NC). Spearman’s rank-order correlation was used for correlations among consumer attributes. Chemical data were averaged across replicates.

The whole dataset was divided into three subsets corresponding to three periods: 2011-2012, 2013-2015 and 2016-2017 since volatile quantification was performed by a different technician in each period and the sugars and acids quantifications were performed three times from frozen samples according to these periods. We observed significant differences in chemical abundance among periods. In order to compensate for technical differences, the normalization method of autoscaling^70^ was conducted for each chemical compound within each period such that periods centered to the same mean and standard deviation. Correlations, clustering and predictive models were built on merged data combining three auto-scaled datasets. Pearson correlations were conducted for all pairs of consumer attributes and chemical compounds.

After false discovery rate (FDR) correction, the correlation matrix was plotted with the “corrplot” package in R for visualization of the relationships. Correlations were filtered with p<0.01 and order of variables was ranked via hierarchical clustering. Non-supervised hierarchical clustering analysis was conducted with the “pvclust” package in R using method.hclust=“ward”. AU (Approximately Unbiased) p-values and BP (Bootstrap Probability) p-values were calculated via multiscale bootstrap resampling to measure the confidence of the clusters^71^. Sensory-chemical network was constructed with the “ggraph” package in R. Only significant correlations were retained after Bonferroni correction to reduce the number of edges for better visualization. Significant relationships between nodes were connected by edges. Kleinberg’s authority centrality scores were calculated for each node to reveal importance of the volatiles in the network.

Partial least square models were built with the “PLS” package in R. Sweetness and consumer liking were regressed on sugars, acids, volatiles and texture liking. Three PLS models corresponded to the three datasets: 2011-2012, 2013-2015 and 2016-2017. The variable importance for the projection (VIP)^72^ was calculated with the R package “PLSVarSel” to reveal important compounds for predictions. We selected 3 principal components for all models due to the smallest square root of mean square error (RMSE) for this number of components (data not shown). A chemical compound was deemed to be important to predict a sensory response if it had a minimum VIP score of 1.0^73^ in two or more of the three datasets. For each of the compounds, a t-test on the slope of the volatile β was applied in a multiple linear regression containing the volatile and total sugars.

Machine learning algorithms were compared for prediction accuracy of liking and sweetness with the “caret” package in R. The whole dataset was split randomly into training (90%) and test (10%) sets for cross-validation (CV). Training dataset were then trained with Random Forest (RF), bayesGLM, GLM, GLMBOOST, LASSO, and PLS models. For comparison, a simple GLM model of individual sugars and acids was constructed using only sucrose, glucose, fructose, malic acid and citric acid. SVMlinear and xgbDART models were also tested but are not reported due to low performance and long computational time, respectively. Nested cross validation of five-fold was performed within 90% training data to tune the hyperparameters. A hundred iterations of nested cross validation were performed to evaluate the stability of model performance.

### Genetic diversity analysis

Twenty-six cultivars and selections included in the sensory study were also genotyped (Table S7). These included UF germplasm, UC-Davis cultivars and two cultivars from other origins. Genotyping was performed using the 38K Axiom® IStraw35 384HT array^37^. Principal component analysis (PCA) was conducted to examine genetic relationships among samples from different origins. SNPs with missing data were omitted in the PCA analyses. A similar PCA was also conducted using chemical data. Individual plots were plotted with the “factoextra” package in R.

### Genetic association analysis

Eight experimental populations, genotyping methods and volatile collection and quantification methods for this analysis are described previously^74^. Relative abundances of individual esters were normalized via the Box-Cox transformation algorithm performed in R-studio prior to genetic correlation. Association analysis was performed using the mixed linear model method implemented in GAPIT v3^75^ in R, using marker positions oriented to the linkage map derived from the ‘Holiday’ x ‘Korona’ mapping population^76^. Genetic associations were evaluated for significance based on the presence of multiple co-locating markers of p-value < 0.05 after FDR multiple comparisons correction.

## Acknowledgments

The authors acknowledge the significant efforts of the UF strawberry breeding program technical staff in growing and maintaining breeding trials, harvests and fruit transportation, as well as the staff members of the Sims and Colquhoun labs in processing and analyzing such large amounts of samples.

## Author contributions

ZF and VW designed the study. ZF and TH performed data analyses. TJ, DG and TC performed chemical analyses. CB performed genetic association analysis. CS organized and performed consumer sensory panels. MR provided direction on data analyses. TC, CS and VW oversaw the whole project. ZF and VW composed the manuscript. All authors read and approved the final manuscript.

## Conflict of Interest

Author Tomas Hasing performed his part of this study while employed at UF/IFAS GCREC and was later employed by Elo Life Systems, Durham, NC, United States by the time of submission. Author Timothy Johnson performed his part of this study while employed at UF/IFAS Environmental Horticulture and was later employed by Driscoll’s Strawberry Associates by the time of submission. The remaining authors declare that the research was conducted in the absence of any commercial or financial relationships that could be construed as a potential conflict of interest.

## Supplementary Information

is available for this paper. Correspondence and requests for materials should be addressed to Zhen Fan at. Reprints and permissions information is available at www.nature.com/reprints

## Extended table legends

Table S1. (A) Correlation matrix among sensory characteristics. (B) Partial correlations among sourness, texture, flavor and liking after controlling for sweetness.

Table S2. Least square means of sensory attributes for each sample.

Table S3. Full chemical data set.

Table S4. Full correlation matrix among sensory attributes, sugars, acids and volatiles.

Table S5. Full partial least square results for sweetness and liking models.

Table S6. Basic sample information.

Table S7. 38K Axiom® IStraw35 384HT array SNP calls of 26 cultivars and selections included in the sensory study.

Figure S1. Manhattan plots of six medium-chain esters.

